# An optimized approach to study sub-sarcomere structure utilizing super-resolution microscopy with secondary VHH nanobodies

**DOI:** 10.1101/2022.09.07.506832

**Authors:** CM Douglas, JE Bird, D Kopinke, KA Esser

## Abstract

The sarcomere is the fundamental contractile unit in skeletal muscle, and the maintenance of its structure is critical for its function. While alterations in sarcomere structure are implicated in many clinical conditions of muscle weakness this area has made limited progress due, in part, to limitations in the ability to robustly detect and measure at sub-sarcomere resolution. Classically the field has relied on approaches including confocal and electron microscopy, but there are technique-specific limitations with respect to resolution, tissue morphology, and protein specific labeling. In this study, our goal was to establish a robust and reproducible method to probe sub-sarcomere protein localization in longitudinal muscle sections. We optimized several steps from tissue preparation to antibody selection and imaging to provide the ability to quantitatively assess spatial distribution of proteins within a single sarcomere. This includes 1) *in situ* fixation for structural integrity, 2) use of multiple same host-species primary antibodies with Fab fragment antibody blocking to maintain specificity, and 3) the use of super-resolution structured illumination microscopy (SIM) to improve from confocal, along with use of emergent VHH secondary nanobodies to double the resolution. The combination of these methods provides a unique approach to improve visualization of sarcomere structure while simultaneously providing the ability to rigorously probe protein localization. While this study focused on assessment of skeletal muscle structure and provides an important set of tools for analysis of skeletal muscle health in disease and aging, we suggest the methods herein may prove advantageous for research outside of skeletal muscle.

## Introduction

Histology—the study of the microscopic structure of tissues—has historically been used to investigate the structure of biological tissues and microorganisms (Mazzarini *et al*., 2021). Tissue structure and the localization of specific proteins can provide insight into the overall function of the tissue or specific protein under investigation. The sarcomere, which is the individual functional unit of skeletal muscle, offers a prime example of this structure-function relationship in biological tissue (Huxley & Hanson, 1954; Huxley & Niedergerke, 1954; Lesanpezeshki *et al*., 2021). Alterations in sarcomere structure have known implications for maximal force-producing capabilities, and emerging data demonstrates the importance of sub-sarcomere structure and protein localization, as well (Lieber & Ward, 2011; Young *et al*., 2017; Hou, 2018; Swist *et al*., 2020). This structure-function relationship of the sarcomere is conserved across species and is important for overall tissue function (Lieber & Ward, 2011; Young et al., 2017; Hou, 2018; Haug et al., 2022). Numerous studies demonstrate this link, but the field has been limited due to the lack of robust methods to quantitatively assess protein localization within the ~2μm sarcomere. Thus, the goal of this study was to develop methods that addressed issues related to these analytical obstacles, including the preservation of structural integrity and incorporation of new imaging tools alongside emergent reagents for sarcomere structural analysis. We propose that these methods will be an important set of tools for analysis of muscle health in diseases and aging.

Aging and diseases such as diabetes, cancer, and Alzheimer’s have been linked to reduced skeletal muscle strength (Frontera *et al*., 2000; Hayes *et al*., 2005; Seene & Kaasik, 2012; Regan *et al*., 2017; Ogawa *et al*., 2018; Moon *et al*., 2019; Huot *et al*., 2021). Further, declining skeletal muscle strength is a major indicator of increased mortality even in healthy, aged individuals (Wolfe, 2006; McGregor *et al*., 2014; García-Hermoso *et al*., 2018; Lesanpezeshki *et al*., 2021). However, these studies largely focus on cross-sectional area as an indicator of overall muscle quality (Frontera *et al*., 2000; Geiger *et al*., 2000; Chimenti *et al*., 2008; Andrews *et al*., 2010; Toth *et al*., 2013; McGregor *et al*., 2014). Although skeletal muscle cross-sectional area positively correlates with overall muscle strength, these measures often do not correlate to muscle quality with respect to sarcomere structure (Frontera *et al*., 2000; Ogawa *et al*., 2018). These studies fail to consider the longitudinal plane of skeletal muscle, where much of the sarcomere is visible. Studies that do include this critical plane of skeletal muscle often rely on measures of sarcomere ultrastructure (i.e. measures of sarcomere length, or global structure) and do not attempt to analyze sub-sarcomere structure (i.e. specific sarcomere protein localization or protein-protein interactions), which is of growing recognition and equal importance to overall sarcomere function (Thompson & Metzger, 2014; Helms *et al*., 2014; Swist *et al*., 2020; Martin *et al*., 2021). Thus, the study of sarcomere structure, and specifically sub-sarcomere structure, is an area of biomedical research with great clinical significance and numerous fundamental questions yet to be answered.

Sarcomere structural analysis has been performed through utilizing various imaging techniques, albeit all with specific pitfalls. The first high-resolution images of sarcomeres were obtained with what many consider the gold standard of the field—electron microscopy (Sjöström & Squire, 1977; Powers *et al*., 2021). This approach provides nanoscopic resolution but is expensive in terms of both sample cost and technical training required, is difficult to label individual proteins, and is notorious for structural artefacts introduced during sample preparation (Ayache *et al*., 2010; Michen *et al*., 2015). Another approach has been the isolation of individual myofibers from muscle tissue with subsequent immunolabeling and imaging (Pasut *et al*., 2013; Horstick *et al*., 2013; Vogler *et al*., 2016; Gallot *et al*., 2016; Lim *et al*., 2018). However, this approach has been acknowledged as relatively harsh and may disrupt structure, it is significantly time-consuming, and many fibers are rejected leading to waste of precious sample tissue (Herzog, 2022). Additionally, sample thickness compared to traditional histological sections can introduce imaging difficulties and various z-plane imaging artefacts (Herzog, 2022). More recently, *in vivo* imaging modalities have been utilized to obtain muscle quality data from sarcomeres *in situ* (Llewellyn *et al*., 2008; Moo *et al*., 2016). However, this approach is label-free and can only provide ultrastructural data compared to both electron microscopy and individual myofibers. These approaches can provide data relating to sarcomere length (Z-disc to Z-disc) and its homogeneity but cannot analyze specific protein localizations or interactions. Finally, traditional approaches of immunohistochemistry with either fresh-frozen cryosections or fresh-frozen paraffin-embedded sections have also been used. However, these approaches have a stringent dichotomy between preserved tissue morphology and antigen epitope availability, due to differences in the tissue preservation processes (Chen *et al*., 2010; Shi *et al*., 2011, 2016; Kumar *et al*., 2015; Hira *et al*., 2019; Liu & Domellöf, 2020). There is a critical need for a robust, reproducible, and technically straightforward approach to study sarcomere sub-structure which can utilize commercially available antibodies and take advantage of developments in high-resolution microscopy, to allow for the specific labeling of multiple proteins, with preserved sarcomere structure.

The primary goal of this study was to develop a robust and reproducible method to preserve sarcomere structure and obtain high-confidence measures of sub-sarcomere structure using longitudinal cryosections. We additionally present a simple and reproducible image processing pipeline – implemented prior to analysis – designed to reduce inter- and intra-study bias and variability in sarcomere structural analysis. The methods introduced herein have great potential for the skeletal muscle field along with additional far-reaching applications for researchers wanting to achieve nanoscopic resolution to study tissue morphology.

## Methods

### Ethical Approval

All animal work performed in this study was conducted in accordance with and approved by the Institutional Animal Care and Use Committee at the University of Florida (IACUC #202009900). The use of animals for all experiments was in accordance with the guidelines established by the US Public Health Service Policy on Humane Care and Use of Laboratory Animals.

### Animals and Tissue Collection

3 male C57BL6 mice (7-10 months of age) were used for all experiments. Mice were housed under a 12:12 light-cycle corresponding to lights on at 7am and lights off at 7pm, with *ad libitum* access to standard rodent chow (Envigo Teklad 2918, Indianapolis, IN, USA) and water. Mice were anaesthetized using isoflurane followed by secondary euthanasia via cervical dislocation as approved by UF IACUC. Tissue collections were performed at a similar time daily (~3pm) to avoid any potential time of day effects in muscle.

### Muscle processing for histological sectioning

Process of muscle preparation for fixation and downstream cryosectioning is seen in Figure 1 (modified from work by (Glancy *et al*., 2015). Entire legs of mice had skin completely removed before being fully removed above the knee. After removal, legs were placed on the flat side of a cork that was cut in half through the middle, lengthwise, with the tibialis anterior (TA) muscle facing upwards away from the cork to better expose it to fixative. The legs were pinned in place with needles through the remaining tissue above the knee, one through the gastrocnemius muscle, and one through the mid-foot with the knee and ankle joints fixed in place at 90° relative to the tibia to standardize muscle length. Corks with leg attached were then placed in a 50mL screw-top conical tube and completely submerged in 10% neutral-buffered formalin (Fisher Healthcare, USA; Cat #23-305510) and left overnight at 4°C. The following day, formalin was replaced with Tris-buffered saline (TBS) solution to wash for 5 minutes. Following this brief wash, the TA muscle is carefully removed from the leg, placed in TBS for an additional 5 minutes before transfer to a 15mL screw-top conical tube filled with 15% sucrose solution (in TBS), and placed overnight at 4°C. The following day, 15% sucrose solution was removed and replaced with 30% sucrose solution, and placed overnight at 4°C. The following day, the TA muscle was placed in TBS for 5 minutes before being placed on a kimwipe to absorb residual moisture. The TA was then placed in Tissue-Tek O.C.T. Compound (O.C.T.; Sakura Finetek, USA; Cat #4583) and frozen in liquid nitrogen-cooled isopentane. Tissue blocks were stored in 5mL screw-top conical tubes at −80°C until needed for sectioning and immunolabeling. TA muscles were sectioned longitudinally at a thickness of 7μm, and sections obtained medially with reference to the longitudinal plane were used for downstream immunolabeling. Tibialis anterior muscles from the contralateral leg were used for control, non-fixed tissues. Control muscle length was standardized to a similar length as fixed tissue by pinning of distal and proximal TA tendons during tissue OCT embedding process.

**Figure 1.**
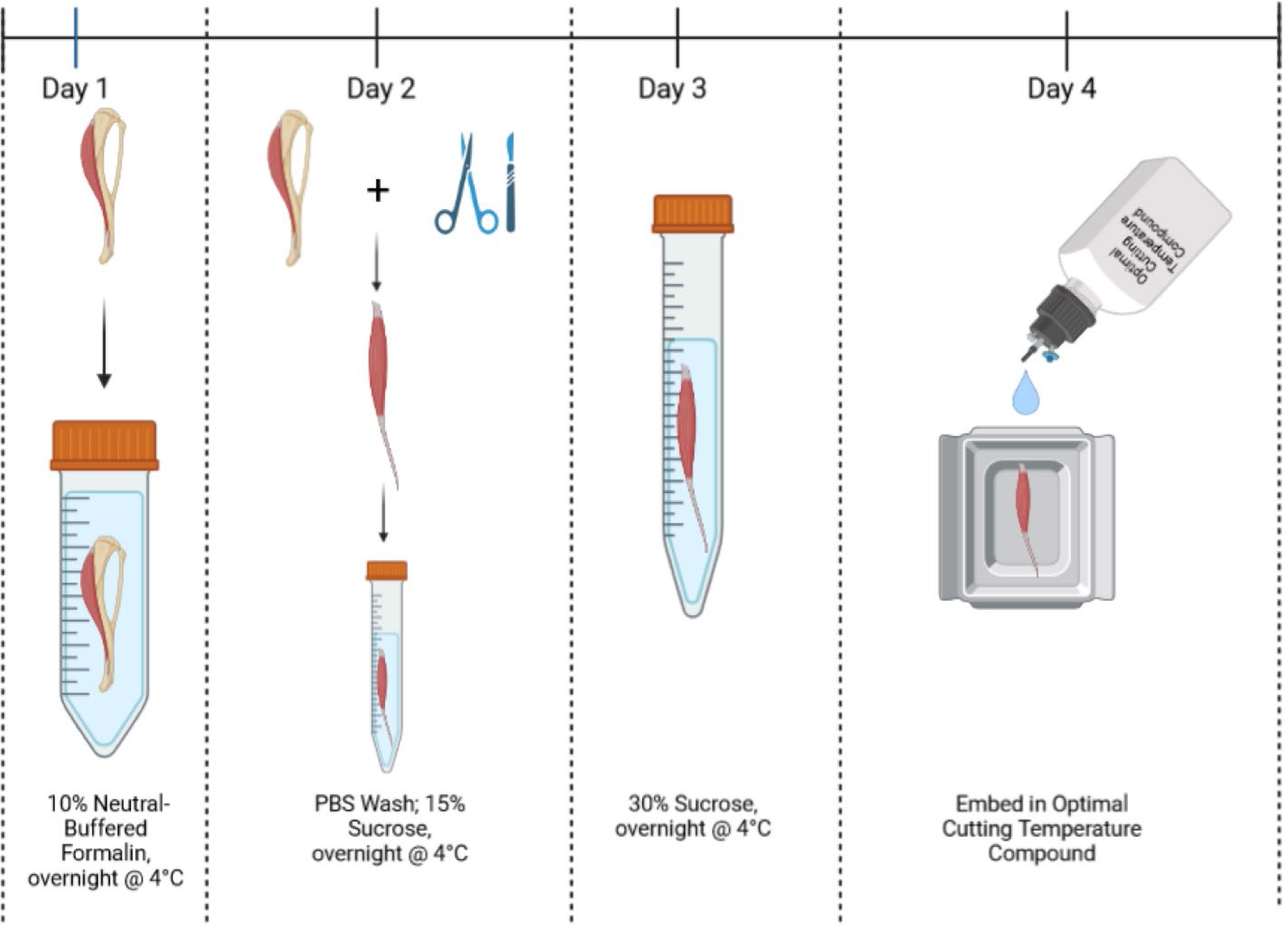
Cartoon demonstrating process of muscle fixation prior to preparation for longitudinal sectioning. In brief, the mouse leg with the tibialis anterior muscle still attached is skinned and removed. The leg is placed on a piece of cork with knee and ankle joints fixed at standardized angles. The leg is then placed in 10% neutral-buffered formalin overnight. The following days, the muscle is removed, washed, and cryoprotected through a sucrose gradient. It is then frozen in OCT compound for cryosectioning. Cartoon generated using BioRender.

### Immunolabeling, Imaging, and antibodies used

An example of the use of multiple same-host primary antibodies labeling scheme is shown in Figure 2, with numbers in black circles showing subsequent steps relating to antibody incubations. Slides were allowed to equilibrate to room temperature (15 minutes) before sections were individually encircled using a hydrophobic PAP pen (Electron Microscopy Sciences, USA; Cat #71312). Sections were then rehydrated for 5 minutes using TBS. Sections were then incubated for 10 minutes with 0.5% Triton X-100 in TBS. Sections were subsequently washed for 5 minutes in TBS, and this wash was repeated 3 times. Following these washes sections were incubated for 30 minutes with one drop of Image-iT™ FX signal enhancer (Invitrogen, USA; Cat #I36933). Sections were washed with TBS for 5 minutes before a 1-hour incubation at room temperature with blocking solution consisting of 5% normal goat serum, 5% normal alpaca serum, 5% bovine serum albumin, 1% glycine, and 0.1% Triton X-100 in TBS. Following incubation with blocking solution, sections were incubated with primary antibody diluted in blocking solution overnight at 4°C. The next day, sections were washed for 5 minutes in TBS + 0.1% Tween-20 (TBS-T) three times, with a following 5-minute wash in TBS before incubation with secondary antibody diluted in blocking solution for 1 hour at room temperature. Following secondary antibody incubation, sections were washed three times for 5 minutes in TBS-T, with a subsequent 5-minute TBS wash. Prior to additional primary antibody incubation, sections were next incubated for one hour at room temperature with excess unconjugated Fab fragment antibodies (Jackson ImmunoResearch, USA) diluted in blocking buffer to block residual binding sites of primary antibodies. Following incubation sections were washed for 5 minutes in TBS-T three times, and subsequently washed once in TBS for 5 minutes. Sections were then briefly incubated (3 minutes) with 4% paraformaldehyde (PFA) solution, and subsequently washed twice in TBS for 5 minutes each. Addition of additional primary and secondary antibodies followed the same steps following the original incubation in blocking solution, for up to two additional primary and secondary antibodies. Following final incubation sections were washed for 5 minutes once in ultrapure water before removal of residual liquid and mounted with ProLong™ Glass antifade mountant (Invitrogen, USA; Cat #P36980) using #1.5H coverslips (ThorLabs; Cat #CG15KH1). Antibodies used in this study along with concentrations and sources used may be found in the table below (Table 1).

**Figure 2.**
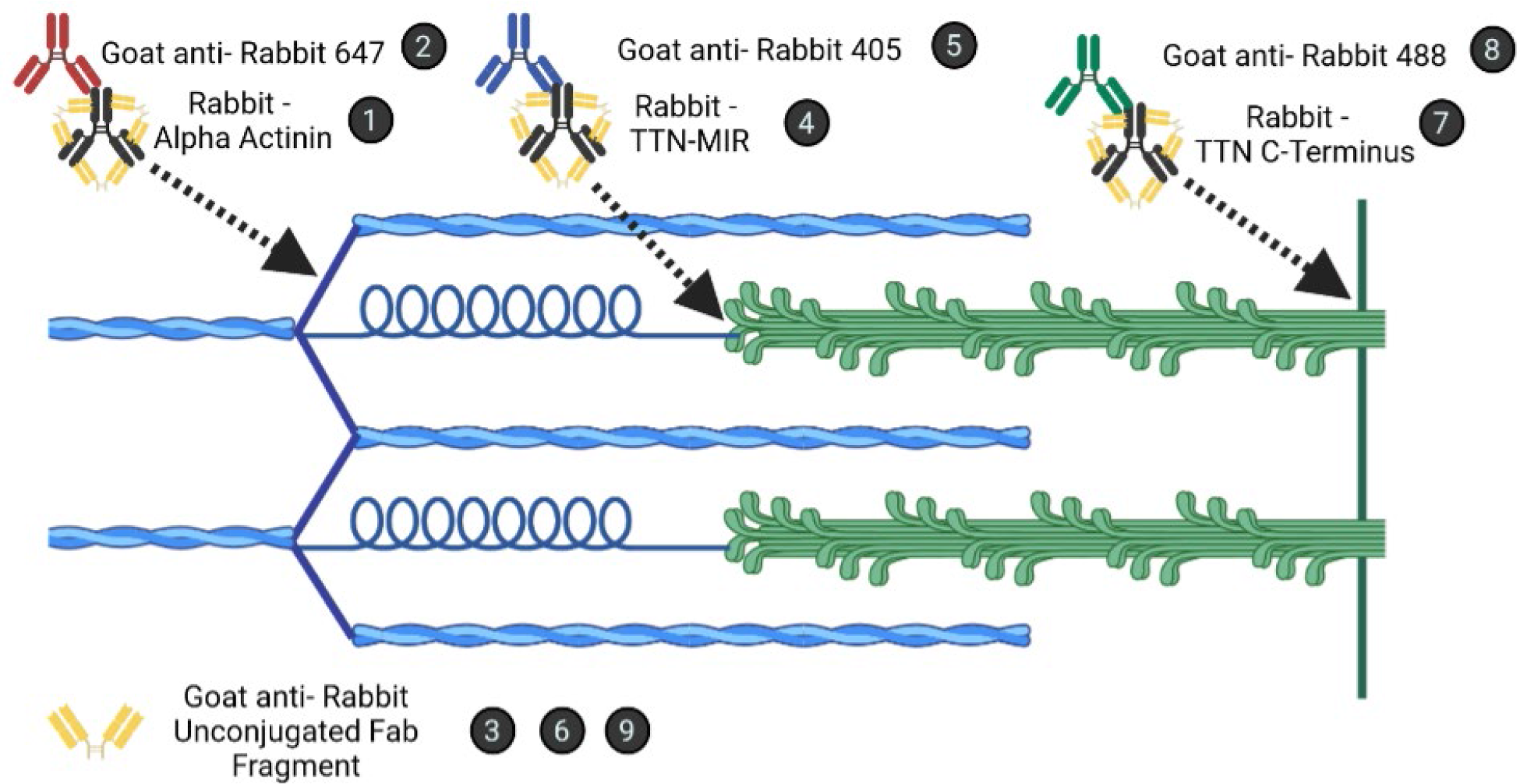
Cartoon (not to scale) demonstrating an example labeling scheme with three same-host primary antibodies with corresponding sarcomere locations across a half-sarcomere (Z-disc to M-line; α-Actinin to TTN C-Terminus). Numbers in black circles correspond to the individual subsequent incubations listed. **1**. α-Actinin antibody is used to label the sarcomere Z-disc. **2**. AlexaFluor 647 secondary antibody is used to label α-Actinin antibody. **3**. Anti-rabbit unconjugated Fab fragment (UFF) is used to block residual open binding sites on α-Actinin primary antibody. **4**. Titin-MIR epitope antibody is used to label the sarcomere A/I-band interface. **5**. AlexaFluor 405 secondary antibody is used to label the Titin-MIR primary antibody. **6**. Anti-rabbit UFF is used to block residual open binding sites on Titin-MIR primary antibody. **7**. Titin C-terminus antibody is used to label the sarcomere M-line, or middle of the sarcomere. **8**. AlexaFluor 488 secondary antibody is used to label Titin C-terminus antibody. **9**. Anti-rabbit UFF is used to block residual open binding sites on Titin C-terminus primary antibody. Cartoon produced with BioRender.

**Table 1.**
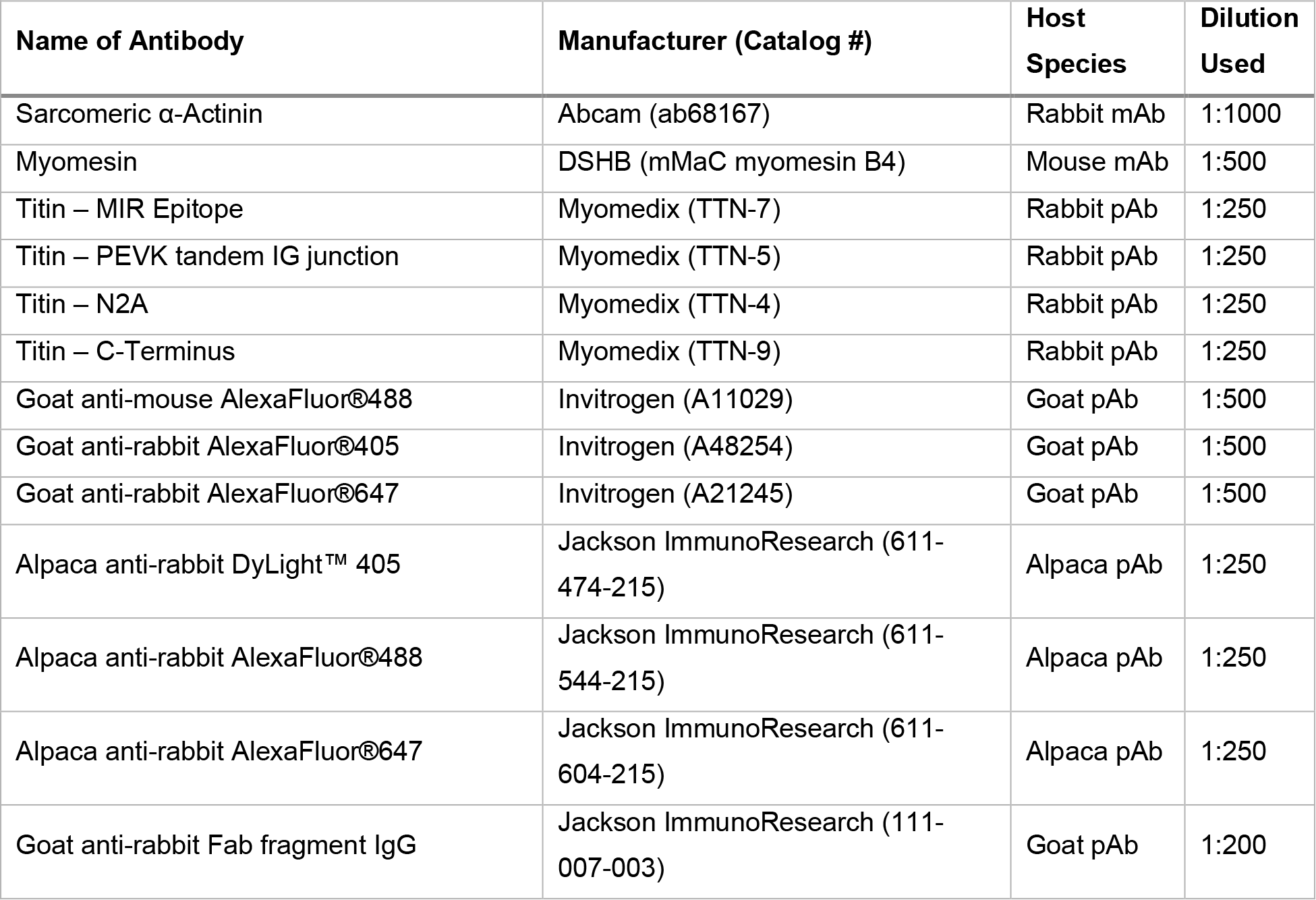
Antibodies.

| Name of Antibody | Manufacturer (Catalog #) | Host Species | Dilution Used |
| --- | --- | --- | --- |
| Sarcomeric α-Actinin | Abcam (ab68167) | Rabbit mAb | 1:1000 |
| Myomesin | DSHB (mMaC myomesin B4) | Mouse mAb | 1:500 |
| Titin – MIR Epitope | Myomedix (TTN-7) | Rabbit pAb | 1:250 |
| Titin – PEVK tandem IG junction | Myomedix (TTN-5) | Rabbit pAb | 1:250 |
| Titin – N2A | Myomedix (TTN-4) | Rabbit pAb | 1:250 |
| Titin – C-Terminus | Myomedix (TTN-9) | Rabbit pAb | 1:250 |
| Goat anti-mouse AlexaFluor®488 | Invitrogen (A11029) | Goat pAb | 1:500 |
| Goat anti-rabbit AlexaFluor®405 | Invitrogen (A48254) | Goat pAb | 1:500 |
| Goat anti-rabbit AlexaFluor®647 | Invitrogen (A21245) | Goat pAb | 1:500 |
| Alpaca anti-rabbit DyLight™ 405 | Jackson ImmunoResearch (611-474-215) | Alpaca pAb | 1:250 |
| Alpaca anti-rabbit AlexaFluor®488 | Jackson ImmunoResearch (611-544-215) | Alpaca pAb | 1:250 |
| Alpaca anti-rabbit AlexaFluor®647 | Jackson ImmunoResearch (611-604-215) | Alpaca pAb | 1:250 |
| Goat anti-rabbit Fab fragment IgG | Jackson ImmunoResearch (111-007-003) | Goat pAb | 1:200 |

Sections were imaged using a Leica SPE Confocal microscope with a separate Leica DFC7000 T Fluorescence high-speed camera under a 63x oil-immersion objective, or with a Nikon Ti2-E inverted microscope equipped with a 100x oil-immersion objective (CFI Apochromat TIRF, NA 1.49) and a confocal spinning disk (Yokogawa, CSU-X1) with super-resolution imaging module (Gataca LiveSR) Systems) and sCMOS camera (Teledyne Photometrics, Prime95B). Z-stacks were obtained in 0.3μm increments set to 300ms exposure per individual channel and reconstructed using maximum intensity slice projections. Image analysis and processing were all performed using ImageJ software. Average background subtraction was performed by use of the rectangle tool to first obtain the mean gray value from a background region of interest outside of the fluorescent label per channel and was subtracted globally from individual channels. Thresholding of individual channels was performed to obtain binary images by use of the *Threshold* function in ImageJ, using the default IsoData algorithm within ImageJ, and applied automatically. Finally, the *Analyze Particles* function was used on individual binary channels to obtain Bare Outline profiles of each binary channel. Individual Bare Outline profiles were then used to obtain distance measures used for sarcomere structural analysis.

### Statistics

All statistical analyses were performed using PRISM software. Statistical comparisons for sarcomere structural measurements (sarcomere length and Z-disc width) are calculated from total measurements per group (n=total measures). Total measurements were derived from 3 biological replicates per group, corresponding to a minimum of 5 non-overlapping line scans performed over each of 5 non-overlapping images per biological replicate for a minimum of 25 line scans per biological replicate. Statistical comparisons between groups of only two was performed using Student’s t-test. Statistical comparisons made between Confocal, SIM, and SIM with VHH nanobodies was performed using one-way ANOVA.

## Results

### Fixation of TA muscles prior to preparation for histological cryosectioning demonstrated better preservation of sarcomere ultrastructure

The first objective was to develop a reproducible protocol to obtain longitudinal cryosections of mouse skeletal muscle tissue with preserved *in vivo* sarcomere ultrastructure. We first developed a process to standardize muscle histological preparations by modification of steps reported by Glancy et al., 2015 (See *Methods*). Once prepared, we obtained longitudinal cryosections from either fixed or contralateral control (non-fixed) tibialis anterior (TA) muscles for visualization and analysis of sarcomere structure via immunofluorescent confocal microscopy. Sections were labeled with antibodies specific to α-Actinin, Myomesin, and the Titin MIR epitope for respective visualization of the Z-disc, M-line, and a point of reference in between. Qualitative and quantitative comparison between fixed and non-fixed TA sections suggested that structure and regularity of all three sarcomere markers were best maintained in fixed TA sections and were representative of sarcomere structure observed via either electron microscopy and *in vivo* imaging modalities (Luther et al., 2002; Llewellyn et al., 2008; Glancy et al., 2014; Moo et al., 2016; Figure 3). Non-fixed TA sections visually demonstrated much less consistency in sarcomere structure compared to fixed TA sections (Figure 3A). Specifically, labeling of α-Actinin and Myomesin suggested more instances of Z-disc streaming and M-line structural heterogeneity in non-fixed TA sections (Figure 3A, bottom) compared to fixed TA sections (Figure 3A, top). These observations suggested that, visually, sarcomere structure was better preserved during our standardized fixation of muscle compared to non-fixed.

**Figure 3.**
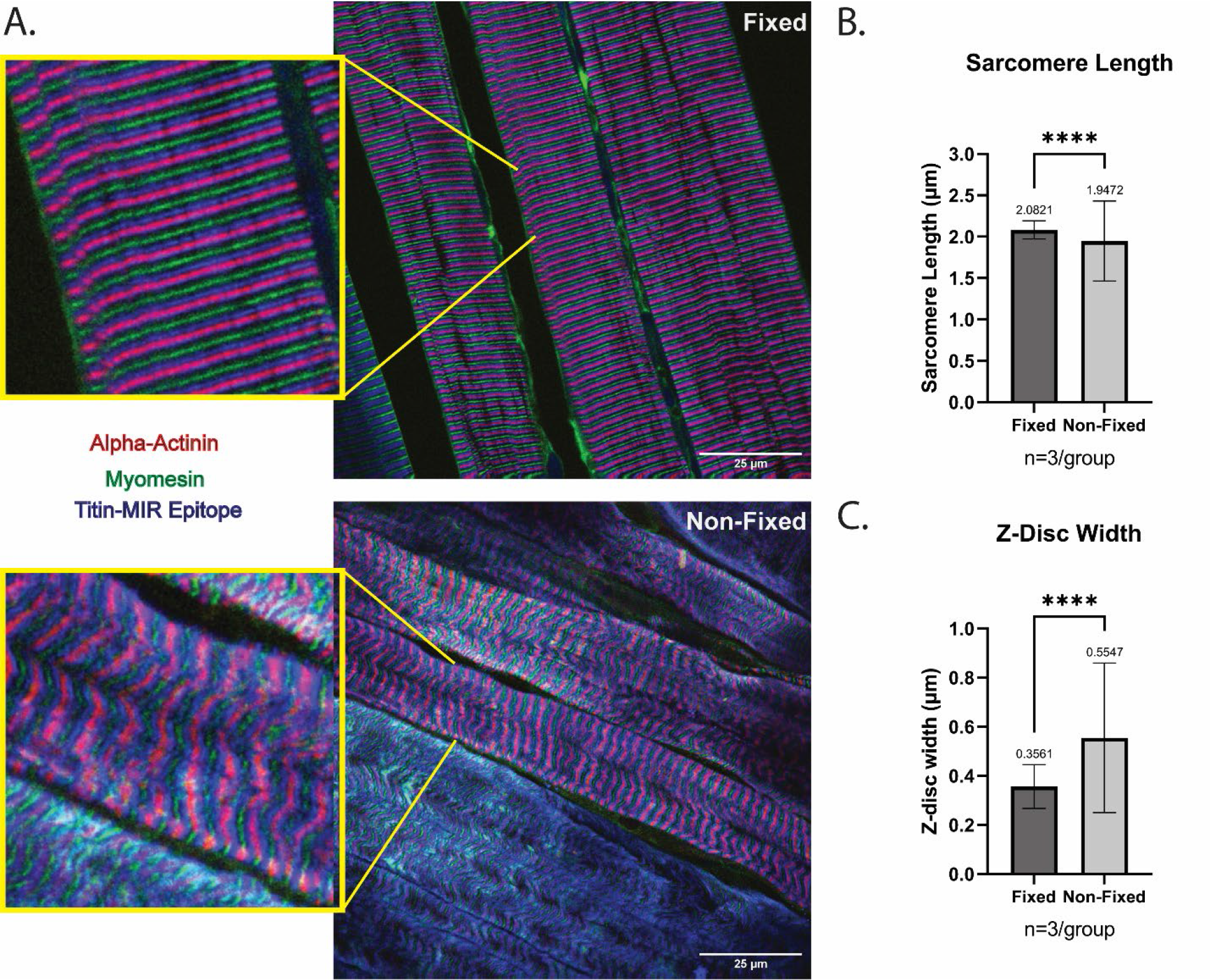
Fixation process prior to embedding muscle for cryosectioning visually and quantitatively preserves sarcomere structure. **A**. Representative images of fixed (top) and non-fixed (bottom) C57BL6 tibialis anterior cryosections labeled with α-Actinin (Z-disc), Myomesin (M-line), and an antibody specific to the Titin-MIR epitope at the sarcomere A/I-band interface. Enhanced images are shown highlighted with yellow border. **B**. Quantification of average sarcomere length between fixed (n=1078) and non-fixed (n=761) TA sections, from centers of neighboring α-Actinin labels. Quantifications are from 5 non-overlapping images acquired from n=3 biological replicates per group. **C**. Quantification of Z-disc width between fixed (n=1191) and non-fixed (n=843) TA sections, and are from 5 non-overlapping images acquired from n=3 biological replicates per group. Plotted measures are shown as mean of total measures with standard deviation, ****p<0.0001.

We next quantified the preservation of *in vivo* sarcomere structure in fixed TA sections compared to non-fixed. Studies using electron microscopy and *in vivo* imaging have demonstrated that homogeneity of both sarcomere length (optimal length ~2.0-2.17; Walker & Schrodt, 1974; Herzog et al., 1992; Gokhin et al., 2014, 2015; Moo et al., 2020) and z-disc width (30-140nm; Luther, 2009; Knöll et al., 2011) are properties of healthy skeletal muscle tissue (Luther et al., 2003; Knöll et al., 2011; Moo et al., 2020). Sarcomere length was obtained by measuring the distance between neighboring centers of α-Actinin labeling; analysis between groups suggested that the average sarcomere length in non-fixed tissue was significantly shorter than fixed tissue (Figure 3B; 1.94±0.017 μm vs. 2.08±0.003 μm; p<0.0001). Importantly, these data also demonstrated that sarcomere length in non-fixed tissue demonstrated less homogeneity than fixed tissue (Figure 3B; F=19.57, p<0.0001). Z-disc width was analyzed by width of α-Actinin fluorescent labeling as the layers of α-Actinin within the Z-disc are commonly used as a reference for Z-disc width (Luther, 2009). Analysis between groups revealed that the average Z-disc width of fixed tissue was significantly narrower compared to non-fixed tissue (0.35±0.002 μm vs. 0.55±0.01 μm; p<0.0001; Figure 3C). Importantly, Z-disc width was significantly less homogenous in non-fixed tissue compared to fixed (F=11.66, p<0.0001; Figure 3C). Overall, these observations indicate that this fixation protocol results in TA cryosections with better preserved ultrastructure with respect to important measures of sarcomere structure.

### Use of post-blocking with unconjugated Fab fragments to allow multiple same-host primary antibody labeling without loss of secondary antibody specificity

We were interested in testing the ability to analyze sub-sarcomere protein associations, but this requires the use of multiple protein-specific primary antibodies on the same section. However, a common issue regarding this approach is that widely available antibodies may be sourced from the same host-species. The resulting potential for non-specific binding of secondary fluorescent antibodies makes image analysis problematic. Recently, canonical IgG antibody fragments retaining the antigen binding domains—Fragment Antigen Binding antibodies, or Fab fragment antibodies—with no additional conjugation have been developed which may be able block residual antigen binding sites allowing for use of multiple same-host primary antibodies (Nelson, 2010). We tested this application to determine if blocking residual binding sites would result in minimal overlap of subsequent secondary antibodies targeting the same host-species primary antibodies (See *Methods*; Figure 2). We utilized three primary antibodies (α-Actinin, the Titin C-terminus, and the Titin-MIR epitope) derived from the same host-species (rabbit) that are specific to three spatially distinct sarcomere regions—the Z-disc, near the M-line, and the sarcomere A/I-band interface, respectively. We first dual-labeled fixed TA sections for both the sarcomere Z-disc and the A/I-band interface, applying primary antibodies (α-Actinin and the Titin-MIR epitope) at the same time and observed the expected overlap of secondary antibodies (Figure 4B). We then applied the primary and secondary antibodies in a sequential order with wash steps in between. Protein labeling was qualitatively more specific, however it was visually apparent that there were still significant instances of overlap between secondary antibodies (Figure 4C). Next, we used the same labeling sequence as before, but applied Fab fragment antibodies after wash steps following the secondary antibodies, to post-block any residual primary antibody antigen-binding sites. We additionally applied a brief incubation with 4% PFA to prevent unbinding of the Fab fragment antibodies from the primary antibodies. We observed little-to-no overlap of secondary antibodies, allowing for definitive localization of primary antibodies from the same host-species (Figure 4D). Finally, we tested the triple-labeling capability of this approach through addition of an M-line specific antibody (Titin C-terminus) (Figure 4A). We observed clear localization of primary antibodies specific to the Z-disc (red), M-line (green), and the A/I-band interface between these two locations (blue) suggesting that post-blocking with Fab fragment antibodies successfully prevents overlap of subsequent secondary antibodies with using same host-species primary antibodies (Figure 4E). These observations demonstrate the use of Fab fragment antibodies as a resource for experiments using at least three same host-species antibodies to study multiple proteins of interest within the same histological section regardless of antibody species origin.

**Figure 4.**
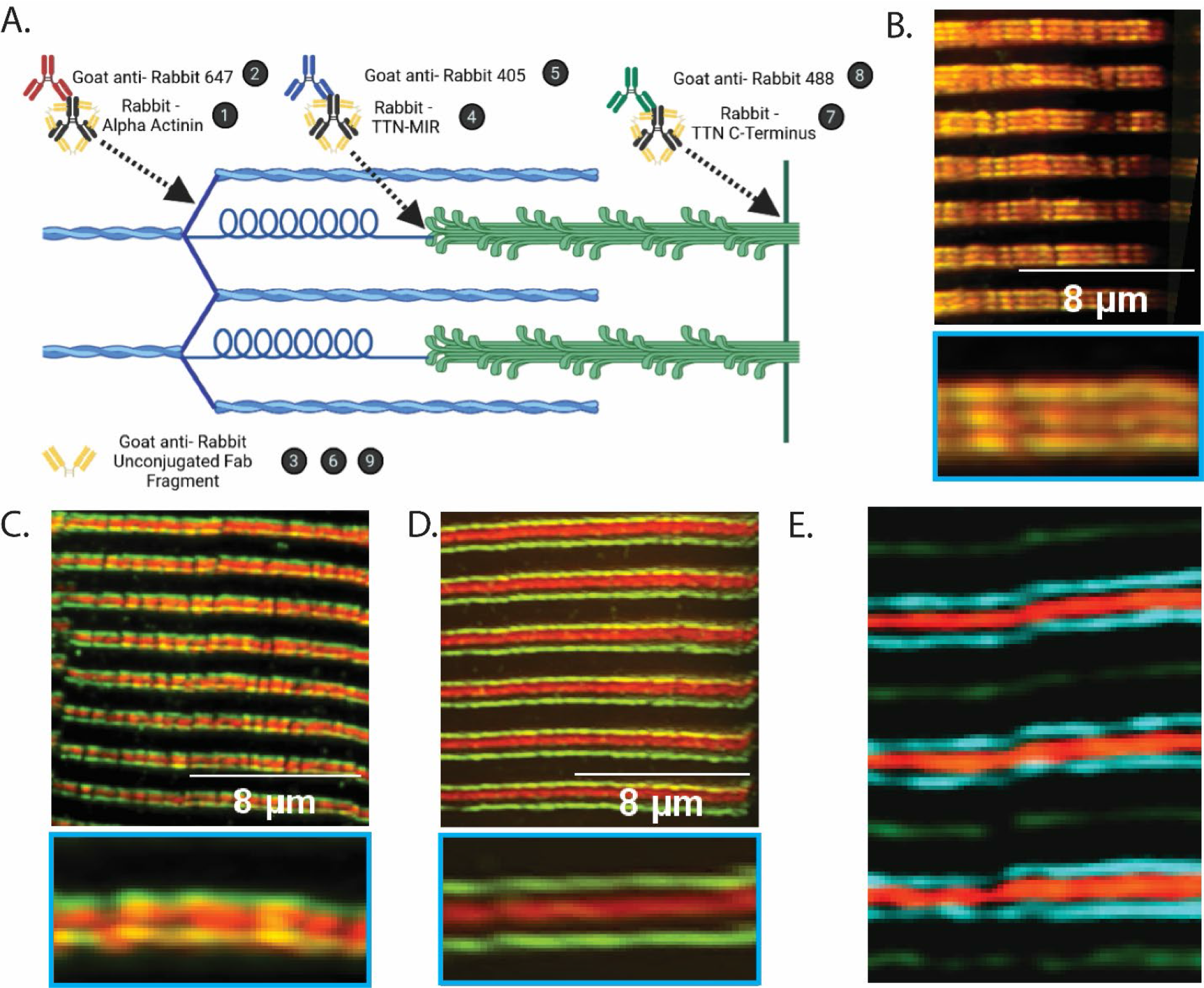
Utilizing a post-blocking step prior to addition of subsequent same-host primary antibodies significantly limits potential overlap of secondary antibody labelings. **A**. Cartoon (produced with BioRender; not to scale) demonstrating an example of the labeling scheme used. Numbers in black circles correspond to individual antibody incubation steps. Panels B, C, and D demonstrate dual labeling using α-Actinin with AlexaFluor 647 conjugation and Titin-MIR with AlexaFluor 488 conjugation. Panel E corresponds to the labeling scheme shown in the cartoon in panel A. Enhanced images of panels B, C, D are highlighted in blue boxes below images. **B**. Representative image showing result of applying primary antibodies at the same time followed by incubation with 488- and 647-conjugated secondary antibodies (Steps 1+4, followed by steps 2+8 in panel A). Secondary antibodies completely overlay which renders protein specificity null. **C**. Representative image demonstrating labeling of primary antibodies in a stepwise manner. Titin-MIR with AlexaFluor 488 added before washing and followed by subsequent addition of α-Actinin and AlexaFluor 647 (Steps 1, 2, wash, 4, 8, in panel A). Overlay of channels indicates dual labeling of secondary antibodies regardless of segmentation of primary antibody steps. **D**. Representative image of labeling with post-blocking steps using anti-Rabbit unconjugated Fab fragments (Step 1, 2, 3, wash, 4, 8, 6 in panel A). Utilization of post-blocking step results in minimal-to-no overlap of secondary antibodies. **E**. Representative image of TA section labeled triple-labeled with same-host primary antibodies from panel A with use of post-blocking Fab fragment antibody step (Step 1, 2, 3, wash, 4, 5, 6, wash, 7, 8, 9 in panel A). Image demonstrates appropriate labeling of secondary antibodies corresponding to three individual antibodies derived from the same host-species (Rabbit).

### Development of an image analysis pipeline to increase accuracy in measures of sarcomere structure

An additional goal was to develop a robust and reproducible image analysis pipeline to reduce both inter- and intra-study variability with respect to measures of sarcomere structure. An approach commonly used in the literature to make measures of protein localization or distance between proteins of interest is through plot profiles derived from line scan analysis. However, this approach is subject to variability or bias based on the point of reference used for protein localization as well as the consistency of the analyst. For example, within the same fluorescent profile plot, the positioning of the line used to measure relative distances among peaks can result in vastly different measures, and this positioning may vary across individual analysts (Figure 5C, top). Further, utilization of common algorithmic processes such as Fourier Transforms to automate these measures are largely optimized for healthy sarcomeres and may inappropriately estimate structure in disordered sarcomeres (Salick *et al*., 2020). We designed a simple and reproducible image pre-processing pipeline within the freely available image analysis software, ImageJ, to reduce the potential for bias and variability in these measures (Figure 5A; see *Methods* for details). In brief, this pipeline consists of:

- An average background subtraction per individual image channels (Figure 5A, left)
- Utilization of the *Threshold* function to binarize the image, promoting foreground and further eliminating background (Figure 5A, center)
- Processing of the binary images using the *Analyze Particles* function to clearly delineate the border of the fluorescent signal (Figure 5A, right)

**Figure 5.**
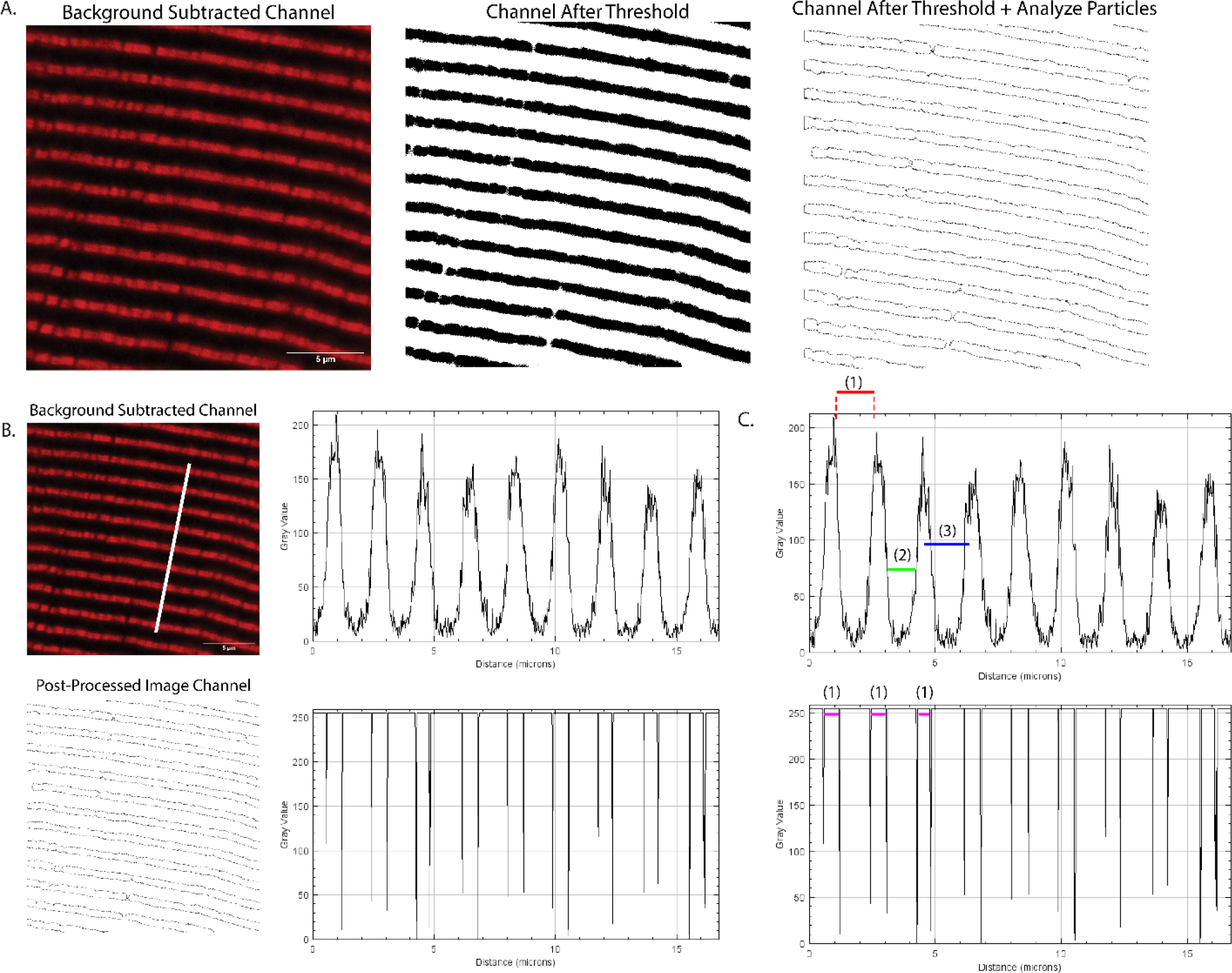
Demonstration of a simple, reproducible approach to generate specified points of reference corresponding to fluorescently labeled cryosections through image pre-processing in ImageJ software. **A**. Example of tibialis anterior cryosection labeled for an antibody specific to the N2A domain of Titin with average background subtracted (left, in red). After using the *Threshold* function, a binarized image remains representing the fluorescent label in black and background in white (middle). Finally, the *Analyze Particles* function in ImageJ processes the binarized image to automatically detect the borders of the fluorescent label (right). **B**. Example images demonstrating the difference in pre- and post-processed images and resulting plot profiles from a line scan (shown as white line, applied across same location in both images). The original image retains vague profile peaks corresponding to fluorescent label (top) while bottom image results in defined lines corresponding to the borders of the fluorescent label (bottom). **C**. Demonstration of differences in points of reference for measures of sarcomere structure between pre- and post-processed images. Without processing, the resulting plot profile allows for multiple points of reference to be used, with variability in distance (points 1, 2, 3; top). Processing of channel results in single, defined points of reference leading to more consistent measures (point 1; bottom).

This pipeline allows for the creation of defined edges of fluorescent labels to be used for points of reference for individual proteins, rather than relying on software to attempt to consistently pick a midpoint from neighboring fluorescent labels. A comparison of the original image used in this example with its corresponding line scan results in a plot profile with undefined, broad peaks (Figure 5B, top) while the processed image results in defined lines indicating the border of the fluorescent label (Figure 5B, bottom). This ultimately results in the elimination of subjectively assigned points of reference (Figure 5C, top) and results in consistently and automatically assigned points of reference (Figure 5C, bottom). By utilizing this pipeline across multiple channels of the same image, measures of multiple sarcomere protein localizations can be performed in a consistent and objective manner.

### Utilizing Super-Resolution microscopy further enhances ability to accurately analyze sarcomere structure

To gain higher resolution for sub-sarcomere analysis, we moved our approach from confocal microscopy to super-resolution structured illumination microscopy (SIM) to 1) test its resolution for accurate measures of Z-disc width and 2) test its ability to delineate specific sarcomere epitopes within nanoscopic proximity. Super-resolution microscopy techniques provide multiple-fold improvements in optical resolution above confocal microscopy (SIM/STED ~2-3 fold; PALM/STORM up to 20-fold), depending on the modality used (Luther, 2009; Galbraith & Galbraith, 2011; Fouquet *et al*., 2015). This increase in resolution allows for more accurate localization of protein epitopes within nanoscopic proximity. For example, fluorescent confocal microscopy cannot distinctly resolve individual localizations of the N2A domain of Titin on either side of the Z-disc (~175nm laterally from the Z-disc; Cazorla et al., 2000), as these are below the diffraction limit. We used this example as a preliminary test of super-resolution microscopy. To resolve the structure of these sub-sarcomere epitopes, we labeled serial fixed TA cryosections with an antibody specific to N2A-Titin and imaged them either with confocal or SIM (Figure 6). We next wanted to test the capacity of this approach to make accurate measures of sub-sarcomere structure and focused on measures of Z-disc width. Measures from studies using electron microscopy report Z-disc width to range from 30-140nm, depending on both fiber type (fast fiber-type are generally more narrow; slow fiber-type are generally wider) and species (Luther, 2009; Knöll *et al*., 2011). The use of fluorescently conjugated antibodies to measure z-disc width has not been achievable due to the resolution limits of confocal microscopy compared to electron microscopy. We therefore believed that performing these measures with SIM would provide the best comparative evidence—with respect to prior electron microscopy measures—to support the robustness of our approach. Analysis of images obtained of fixed TA cryosections labeled for α-Actinin and imaged using either confocal microscopy or SIM demonstrated that Z-disc width obtained using SIM was significantly narrower compared to width measures from images obtained using confocal microscopy (383nm vs. 144nm, p<0.0001; Figure 7A,B,D). Importantly, these measures are much closer to reported measures of Z-disc width with electron microscopy (Luther, 2009; Knöll *et al*., 2011).

**Figure 6.**
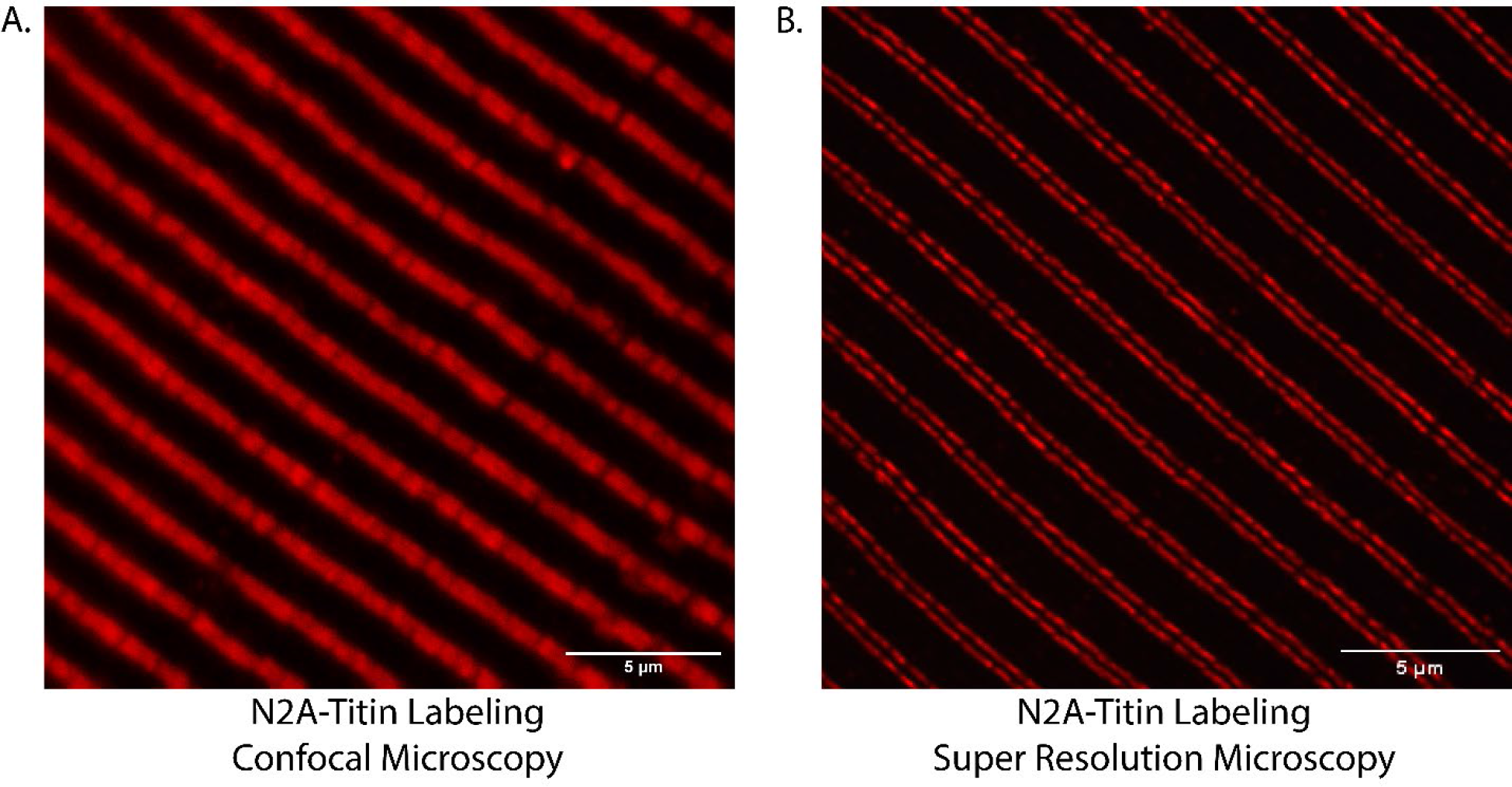
Representative images demonstrating increased resolving capability between confocal and SIM of N2A epitope of Titin in tibialis anterior cryosections. **A**. Image of Titin-N2A labeling imaged using confocal microscopy. Imaging resolves labeling as an individual band. **B**. Image of Titin-N2A labeling imaged with SIM. Imaging resolves labeling as expected doublet pattern on either side of the sarcomere Z-disc.

**Figure 7.**
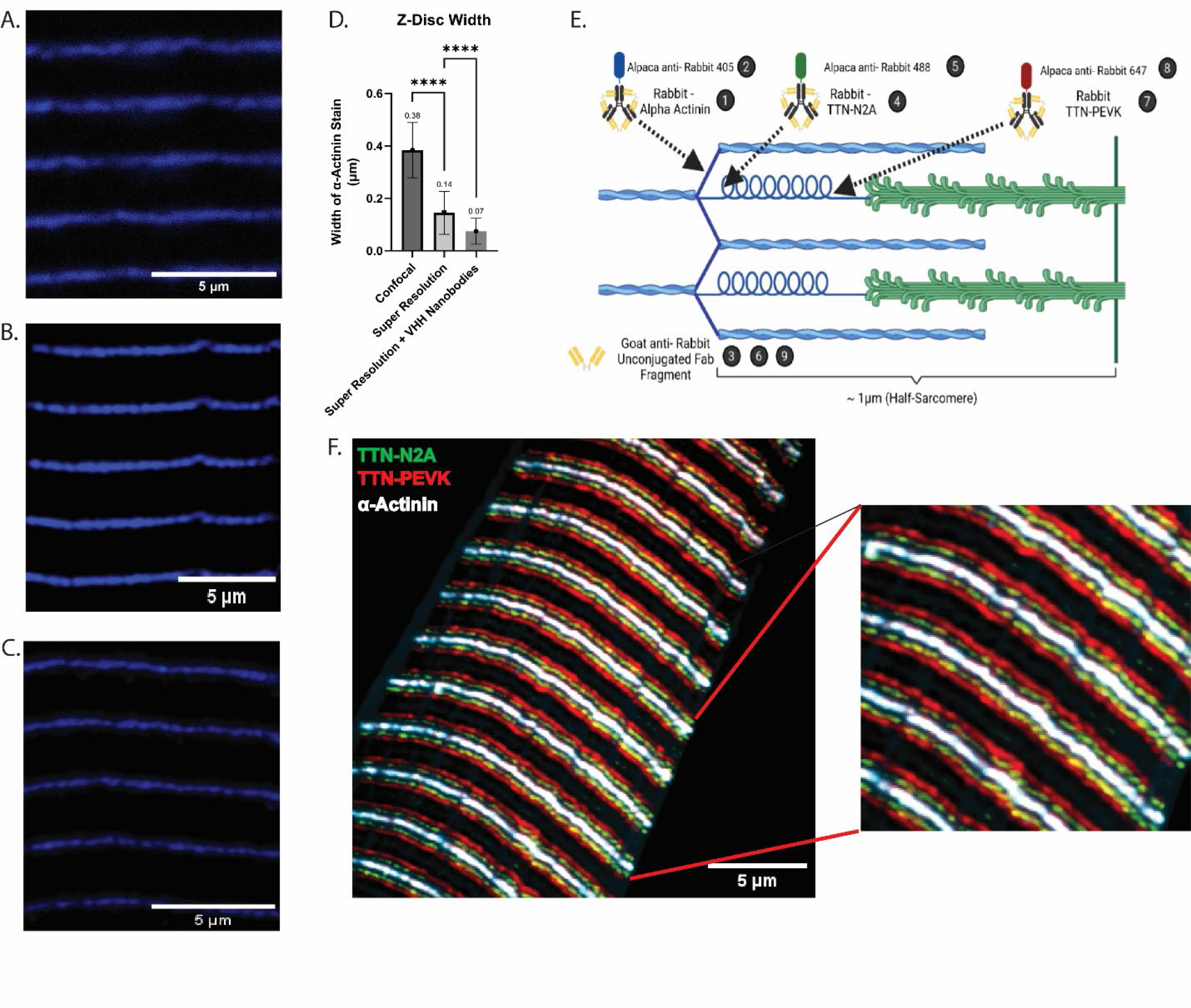
Utilization of SIM in concert with nanobodies further enhances resolution and provides support for preservation of *in vivo* sarcomere structure. **A**. Representative image of α-Actinin labeling with canonical IgG antibodies on tibialis anterior cryosection and imaged using confocal microscopy. **B**. Representative image of α-Actinin labeling with canonical IgG antibodies on tibialis anterior cryosection and imaged using SIM. Borders of labeling are notably more defined than with confocal microscopy. **C**. Representative image of α-Actinin labeling with canonical IgG primary antibodies + VHH-conjugated secondary nanobodies on tibialis anterior cryosection and imaged with SIM. Fluorescent label is notably narrower than SIM without nanobodies. **D**. Quantification of Z-disc width measures obtained from confocal microscopy (n=649), SIM (n=519), and SIM + secondary nanobodies (n=278). Quantifications were performed using n=3 biological replicates per group, and 5 non-overlapping images per biological replicate. **E**. Cartoon (not to scale) depicting labeling scheme used for images seen in panel F. Numbers in black circles represent individual incubations used to perform triple-labeling with same host-species primary antibodies. Cartoon produced using BioRender. **F**. Image demonstrating ability to delineate three sarcomere epitopes of close proximity (N2A-PEVK distance ~100nm) with increased resolution using secondary nanobodies. Tibialis anterior cryosection labeled for α-Actinin (white), Titin-N2A (green), and Titin-PEVK (red) demonstrates excellent resolving capability with preserved sarcomere structure.

Another limitation of immunofluorescent microscopy is the molecular localization of the fluorophore used for visualization with respect to the localization of the protein itself, or linkage error. Traditionally, two IgG antibodies—one primary and one secondary—are needed to label and visualize the protein. This results in a considerable gap at the molecular level between the protein of interest and the fluorophore (~25nm), and thereby limits resolving power upstream of the microscope (Sograte-Idrissi *et al*., 2019). However, by using smaller molecules to label and visualize proteins, some of this linkage error can be reduced. One of the commercial options is the use of newly developed VHH secondary antibodies—or “nanobodies”—produced in alpaca that are much smaller than canonical IgG secondary antibodies (10-fold smaller; 15kDa vs. 150kDa) and reduces this linkage error significantly (from ~25nm to ~4nm) (Desmyter *et al*., 2015; Beghein & Gettemans, 2017; Arbabi-Ghahroudi, 2017; Pleiner *et al*., 2018; Carrington *et al*., 2019; Asaadi *et al*., 2021). We labeled fixed TA cryosections with primary IgG antibodies specific to α-Actinin followed by secondary fluorophore-conjugated VHH nanobodies (Figure 7C, E). We then measured and compared Z-disc width measures obtained with SIM and VHH secondary nanobodies to both confocal and SIM with traditional IgG secondary antibodies (Figure 7D). Imaging with increased resolution with SIM compared to confocal, there was an expected significant increase in the accuracy of Z-disc width (380nm vs. 74nm, p<0.0001; Figure 7D); yet more importantly, comparison of Z-disc width using SIM with nanobodies to SIM with traditional IgG antibodies resulted in a ~100% decrease in width measures of the Z-disc (74nm vs. 144nm, p<0.0001; Figure 7D). Importantly, the addition of secondary nanobodies resulted in measures of Z-disc width that are more aligned with established electron microscopy measures of the Z-disc in fast-twitch muscle fibers (74nm vs. 30-50nm; Luther & Squire, 2002; Luther et al., 2003). We speculate that the addition of primary nanobodies in concert with secondary nanobodies will result in enhanced resolving capacity, and measures even more in line with reported measures from studies using electron microscopy; however, commercially available protein VHH primary nanobodies remain limited to date and will require validation of specificity in future studies.

Lastly, we provide proof of concept that this protocol is sufficient to resolve multiple sub-sarcomere epitopes of interest using our multiple same-host antibody labeling approach in combination with secondary nanobodies. We labeled fixed TA cryosections with antibodies specific to α-Actinin, the N2A domain of Titin, and the PEVK domain of Titin, all of which lie in proximity along the sarcomere (<~200nm total distance, ~100nm from N2A to PEVK Titin epitopes; (Cazorla *et al*., 2000; van der Pijl *et al.*, 2021*a*, 2021*b*); Figure 7E). Images obtained using SIM demonstrate the clear separation of individual epitopes. This confirmed the ability to individually resolve each of these antibodies with minimal-to-no overlap, demonstrating the ability to make measures across different epitopes within a sarcomere at nanoscopic precision (Figure 7F). By increasing the resolving power through biochemical and technical approaches, we demonstrate that our protocol is robust, reproducible, and capable of measuring sub-sarcomere structure well below the micron scale.

## Discussion

The homogeneity of healthy sarcomere structure is important for maintaining the force generating function of muscle tissue. Thus, researchers often turn to the study of muscle morphology for understanding health status of skeletal muscle across various tissue states (i.e., injury, disease, aging). Approaches used to visualize sarcomere structure include light microscopy, electron microscopy, immunohistochemistry, and immunofluorescence with tissue sections or isolated myofibers, as well as *in vivo* imaging modalities. These approaches all have unique limitations, but the most delineating factors between the *ex vivo* and *in vivo* approaches have largely been 1) the preservation of *in situ* morphology, 2) the ability to label and visualize specific proteins, and 3) the imaging resolution provided by each approach. For example, *in vivo* imaging provides details of true representative tissue morphology while *ex vivo* approaches may alter tissue morphology during the tissue preparation process. Additionally, while tissue morphology is in its purest form *in vivo*, these approaches are severely limited by the inability to label and visualize specific proteins compared to *ex vivo* approaches. Finally, while *in vivo* approaches provide details of true tissue morphology, the technical limitations of these approaches cannot provide the level of sub-micron resolution capable of advanced microscopy techniques. There is a need for an approach capable of obtaining nanoscopic resolution with the ability to specifically label proteins or epitopes of interest to further probe elements within sarcomere structure. Therefore, in this study we developed a robust and reproducible protocol to analyze sub-sarcomere structure through use of longitudinal cryosections with preserved *in vivo* morphology along with super-resolution microscopy in concert with emerging fluorescent nanobody technology.

In this study we demonstrate the robustness of our protocol through a transition from confocal to SIM with inclusion of emergent nanobody technology. This combination allows us to use immunofluorescence to accurately measure Z-disc widths like those reported using electron microscopy. In addition, we can resolve distances between proteins or specific epitopes as close as ~100nm and probe the localization of up to three targeted antibodies of the same host-species. We suggest that use of this approach with models of muscle weakness holds potential to help identify potential changes in protein localization and potential disruptions in sub-sarcomere structure that could contribute to functional declines. We note that our approach is not the first to utilize advanced imaging modalities to study skeletal muscle structure (Radermacher *et al*., 1994; Blanc *et al*., 2000; Soeller & Baddeley, 2013; Glancy *et al*., 2014; Sun *et al*., 2014; Jayasinghe *et al*., 2014, 2015, 2018; Agarwal & Machá, 2016; York & Zheng, 2017; Gemmink *et al*., 2018; Yi *et al*., 2019; Ghosh *et al*., 2019; Szikora *et al*., 2020; Daneshparvar *et al*., 2020; Oda & Yanagisawa, 2020; Hofemeier *et al*., 2021; Rahmanseresht *et al*., 2021; van der Pijl *et al*., 2021*a*; Wang *et al*., 2021; Melville *et al*., 2022; Rimoli *et al*., 2022; Skorska *et al*., 2022; Schueder *et al*., 2022; McMillan & Scarff, 2022; Shaib *et al.*, 2022). Many of these approaches implement single molecule super-resolution microscopy techniques, such as stochastic optical reconstruction microscopy (STORM). These approaches, while providing greater localization accuracy compared to SIM, require comparatively extensive experimental set-up, specialized fluorophores, more time required per image, and extensive training required for optimal results. We believe that our protocol with fluorescent nanobodies in combination with SIM provides an advantage in terms of less technical experience required and speed of image capture (seconds vs. minutes). In particular, the use of emerging nanobody technology is exciting. Although discovered over 25 years ago, these VHH nanobodies are now being used to create conjugated antibodies of much smaller mass (~15 kDa) compared to traditionally used whole molecule IgG (~150 kDa). The use of these secondary nanobodies eliminates a considerable portion of the linkage error that commonly plagues protein localization using immunofluorescence microscopy. In addition, these nanobodies in theory will allow for better penetrance in tissue sections and may more effectively label protein epitopes in fixed tissues that canonical IgG antibodies can no longer recognize. To our knowledge, our protocol is the first to demonstrate the usage of secondary nanobodies to improve accuracy of protein localization in skeletal muscle using SIM, and we suggest that future studies take advantage of this emergent technology to improve protein localization accuracy regardless of imaging modality.

For skeletal muscle research, we believe that our protocol will help support more studies focused on the study of sarcomere sub-structure. At the molecular level, individual sarcomeres (~2.0-2.17 μm in length) are comparatively large structures composed of dozens of proteins; many of these proteins reside within distinct sub-regions, such as the Z-disc or M-line. Prior to newer approaches, the resolution of protein localization within the sarcomere has been limited due to diffraction limited imaging techniques. As such, many studies have historically referenced specific protein localizations to general sarcomere regions—i.e., the Z-disc, M-line, etc. Yet within these large protein complexes, there are nuances to specific protein localizations and interactions. There is growing recognition of the importance for their individual roles within these sub-sarcomere domains such as modulation of muscle function, signaling, or overall sarcomere structure. With advances in protein localization resolving power due to emergent nanobodies, studies can confidently begin to visualize the nanoscopic specificity of protein localization within these sub-regions (i.e., central, or distal to the Z-disc center), or with respect to other proteins (i.e., localization of proteins relative to a specific epitope of Titin, or specifically where Titin’s N2A epitope interacts with the Actin filament, etc.). Further, utilization of primary nanobodies—alongside secondary nanobodies— when they become commercially available, should act to further enhance the resolving power of fluorescence microscopy by shrinking the molecular gap between protein and fluorophore. We encourage future studies to take advantage of these recent developments, as well as future improvements in antibody technology to better demonstrate the research potential of these non-canonical antibodies.

## Conclusions

The study of sarcomere structure is critical in understanding the overall health of the tissue. Most studies of muscle health and disease focus on cross-sectional area, but fewer studies have focused on longitudinal structure—where sarcomeres are readily visible—due to technical challenges. Additionally, there is a growing recognition for the contributions of proteins with specific sub-sarcomere localizations such as the M-line, Z-disc, or along Titin filaments, as modulators of muscle function, signaling, and structure. Therefore, there is a need for an approach capable of obtaining nanoscopic resolution of *in vivo* sarcomere morphology with the capacity to label specific proteins of interest to better understand structure. We present a simple, robust, and reproducible method to obtain and analyze longitudinal sections of mouse TA muscle with preserved structure that is representative of *in vivo* sarcomere structure. We provide instructions for labeling multiple sarcomere proteins using a scheme not limited by host species of the primary antibodies. We further provide examples of using newly emerging VHH-nanobody technology to further enhance resolving power with advanced imaging modalities and demonstrate the potential for assessing sub-sarcomere structure and protein localization. These methods presented in this study will assist not only the skeletal muscle field in the understanding of sarcomere structure but will allow others to study their tissue of interest at the nanoscopic level.

## Acknowledgements

We thank Alessandra Norris (University of Florida) for technical assistance with confocal microscopy experiments. This work was supported by National Institutes of Health grant 1R01AR07922001 to KAE, 1R01AR079449 to DK, as well as the University of Florida.

## Notes

### Competing Interest Statement

The authors have declared no competing interest.

